# Transcriptomic gene profiles in an *ex vivo* model of erythropoiesis to unravel molecular pathomechanisms of Sickle Cell Disease

**DOI:** 10.1101/2023.09.26.559266

**Authors:** Matthis Tinguely, Lennart Opitz, Dominik J. Schaer, Florence Vallelian, Markus Schmugge, Francesca D. Franzoso

## Abstract

We characterized the transcriptional profiles of erythroid cells differentiated from peripheral blood mononuclear cells (PBMCs) from peripheral blood collected from patients diagnosed with Sickle Cell Disease (SCD), which have been treated with Hydroxyurea (HU) in comparison to untreated SCD patients and healthy controls (HC) using bulk RNAseq. We identified 1398 differentially expressed genes (DEGs) in SCD non-treated-derived erythroid cells and 495 DEGs in SCD HU-treated patient-derived erythroid cells compared to HC. We found biological processes such as oxidative phosphorylation pathway, proteasome, autophagy, natural killer cell (NK) cytotoxicity, adaptive immune response or inflammatory response to be significantly enriched in our patient study groups by using Gene Ontology (GO) and Kyoto Encyclopedia of Genes and Genomes (KEGG) analysis. Our findings collectively suggest different as well as common molecular signatures between our groups. We could validate 12 of our top DEGs in treated patients by qRT-PCR. We found similar regulation patterns when comparing the mRNA levels of mutS homolog 5-Suppressor APC Domain Containing 1 (MSH5-SAPCD1), G protein subunit gamma 4 (GNG4), stabilin 1/ clever-1 (STAB1) and Fas Binding Factor 1 (FBF1) from the bone marrow cells and spleen tissue from the Berkely SCD mouse model to the expressions observed in the transcriptome of our ex-vivo patient-derived erythropoiesis model.

## Introduction

Sickle cell disease (SCD) is an autosomal recessive, monogenetic disorder.^1^ Sickled haemoglobin (HbS - α_2_β^S^_2_) was described as a structural variant of normal adult haemoglobin (HbA) caused by a point mutation in the *HBB* gene that leads to the single amino acid substitution of valine for glutamic acid at position 6 of the β-globin subunit (β^S^) of the haemoglobin molecule.^1–3^ Hydroxyurea (HU), a ribonucleoside diphosphate reductase inhibitor, originally used to reduce the need of blood transfusions and the frequency of 3 vaso-occlusive painful crisis each year in adults with homozygous SCD,^4^ has remained as the first-line therapy. ^5^The mechanisms of HU in SCD primarily involves increasing production of fetal haemoglobin (HbF).^6^ HU treatment in adult patients for up to 9 years were associated with a significant reduction of 40% in mortality.^75^

Several studies have previously analysed transcriptional signalling pathways and differentially expressed genes (DEGs) that may be dysregulated in SCD. Raghavachari et al.^8^ identified in a platelet transcriptome analysis in SCD patients a range of multiple platelet-specific genes involved in arginine metabolism and redox homeostasis. Top platelet-abundant genes included platelet derived growth factor receptor alpha (PDGFRA), myosin light chain kinase (MYLK), platelet factor 4 variant 1 (PF4V1), 5 ‘-aminolevulinate synthase 2 (ALAS2), histone cluster 1, H2bg (HIST1H2BG) and solute carrier family 24 member 3 (SLC24A3).^8^ Creary et al.^9^ found changes in gene expression profiles using whole blood RNA-Sequencing among patients with SCD hospitalized for acute chest syndrome (ACS) and acute vaso-occlusive pain crises (VOC) episodes. Top DEGs included CD177 molecule (CD177), caspase 5 (CASP5), suppressor of cytokine signalling 3 (SOCS3), free fatty acid receptor 3 (FFAR3), GRAM domain containing 1C (GRAMD1C) and annexin A3 (ANXA3), whereas top canonical pathways identified in ACS were interferon signalling, pattern recognition receptors, neuroinflammation, macrophages and in VOC were related to IL-10 signalling, iNOS signalling, IL-6 signalling and B cell signalling.^9^ The genome-wide gene expression profile in peripheral blood mononuclear cell (PBMC), performed by Desai et al^10^ in 3 cohorts of SCD patients revealed the role of haemolysis and globin synthesis in SCD through gene expression upregulation in BCL2 like 1 (BCL2L1), DOCK family candidates, selenium binding protein 1 (SELENBP1), Delta-aminolevulinate synthase 2 (ALAS2), basigin (BSG), ferrochelatase (FECH), glycophorin A (GYPA), and bisphosphoglycerate mutase (BPGM), as well as upregulation of different metabolic pathways, complement and coagulation cascades, malaria signalling and downregulation of T-cell receptor signalling, immunodeficiency, and vascular endothelial factor (VEGF) pathway.^10^

Transgenic mouse models and erythroid cell lines as *in vitro* models have been used to understand the underlying mechanisms of pathogenesis of various diseases and as preclinical testing for new therapies. In our study we used an erythropoiesis model using a 2-phase liquid system in which we previously showed that microRNA-96 directly supress γ-globin expression by contributing to HbF regulation.^11,12^ We used the Berkeley SCD transgenic mouse model which includes genetic disruptions of endogenous adult-type α and β-globin gene (*Hbb-a1, Hbb-a2, Hbb-b*^*maj*^ and *Hbb-b*^*min*^) and expresses human globins via three DNA transgenes: 1.5 kb spanning the α-globin gene *HBA1*; a contiguous 39 kb genomic fragment including genes for γ-globin (*HBG2, HBG1*), δ-globin (*HBD*) and sickle β-globin (*HBB*^*S*^); and a 6.5 kb *‘*mini-locus control region (LCR) ‘ derived from an endogenous enhancer in the human β-like globin cluster that confers high-level erythroid expression.^13,14^ Berkeley mice have many similar pathologies as the human SCD^15^ and they have been used to test lentiviral or Cas9-induced DNA repair vectors for beta-globin gene replacement or to alter the mutant SCD codon.^16–21^

In this present study we aimed to further identify the erythroid-specific transcriptional profiles of non-treated and HU-treated SCD patients, in comparison to HC using a patient-derived erythropoiesis *in vitro* model. In our primary patient-derived erythroid cultures we could validate top candidate genes by RT-qPCR and further explore their expression in the bone marrow and spleen of the Berkeley SCD mouse model. The identified genes and pathways could be further considered as important role-players in SCD pathogenesis and might lead to discovering new treatment targets.

## Results

### Participant enrolment

In total, we recruited 17 children with median age: 9.0 ± 5.36 years; gender ratio: 8 females (F) / 9 males (M)). The 5 SCD non-treated patients presented a median age: 10.0 ± 6.05 years; gender ratio: 5F/0M. The 7 SCD HU-treated patients presented a median age: 12.0 ± 6.8 years; gender ratio: 1F/6M). The 5 healthy paediatric controls (HC) had a median age of 8.0 ± 3.28 years (2F/ 3M). All the SCD patients were homozygous for the HbSS genotype. The age of the HC was lower than the SCD patients, p=0.004, Mann Whitney U test. Exclusion criteria were: acute or chronic disease and chronic organ dysfunction apart from SCD itself (i.e., sickle nephropathy, moyamoya disease, iron overload) and history of cardiovascular disease.

### Transcriptome profiling in untreated SCD and HU-treated SCD patients versus healthy controls

We identified by RNA sequencing 1398 DEGs in SCD non-treated derived erythroid cells compared to HC, of which 519 genes were upregulated (e.g., Calneuron 1 (CALN1)) and 879 genes (e.g., Glutathione S-Transferase Theta 2B (GSTT2B), mutS homolog 5-Suppressor APC Domain Containing 1 (MSH5-SAPCD1) and Retinol Dehydrogenase 10 (RDH10)) were downregulated (Figure 1A). 495 DEGs were identified in SCD HU-treated patient-derived erythroid cells compared to HC, of which 288 genes were upregulated (e.g., CALN1) and 207 genes (e.g., GSTT2B, Fas Binding Factor 1 (FBF1) and MSH5-SAPCD1) were downregulated (Figure 1B). 786 DEGs were identified in SCD HU-treated patient-derived erythroid cells compared to SCD non-treated derived erythroid cells, of which 455 genes (e.g., G Protein Subunit Gamma 4 (GNG4) and RDH10)) were upregulated and 331 genes were downregulated (Figure 1C).

We generated heatmaps to display the level of expression of each gene in relation to the mean level of expression of that gene across all samples. Overall, the HU treatment altered the SCD gene expression signatures, some even being reversed to the control levels (Figure 2). Specifically, when we compared SCD non-treated to the HU-treated and HC samples, we could observe a similar upregulation of DEGs in contrast to the downregulated DEGs of non-treated samples (Figure 2A). Whereas, in HU-treated SCD versus HC, we found an overall downregulation of gene expression in HC, compared to upregulated DEGs of non-treated and HU-treated SCD samples for unc-51 like kinase 3 (ULK3), Kinesin Family Member C2 (KIFC2), translocase of inner mitochondrial membrane 8A (TIMM8A), AC004233.2, CU638689.4, AL035461.2, crystallin beta B2 (CRYBB2), coiled-coil glutamate rich protein 2 (CCER2), SLX1 Homolog A, Structure-Specific Endonuclease Subunit (SLX1A), MIR503HG, GSTT2B, Phenylethanolamine N-Methyltransferase (PNMT), AC004832.3, solute carrier 11a1 (SLC11A1), ribonuclease A family member 1 (RNASE1) and desmin (DES). Moreover, genes such as transmembrane 4 L six family member 1 (TM4SF1), CALN1, ADGRA2, long intergenic non-protein coding RNA 2099 (LINC02099), endoplasmic reticulum aminopeptidase 2 (ERAP2), neuron navigator 3 (NAV3), neuregulin 1 (NRG1), solute carrier family 44 member (SLC44A5), AC083862.3, leukocyte immunoglobulin like receptor A4 (LILRA4), ras related glycolysis inhibitor and calcium channel regulator (RRAD), STEAP4 metalloreductase (STEAP4), caveolin 1 (CAV1) and phosphate regulating endopeptidase X-Linked (PHEX), showed a significant upregulation in HC, and a significant downregulation in non-treated and HU-treated SCD samples (Figure 2B). Furthermore, when we compared the expression level of 20 most significant DEGs in SCD HU-treated versus SCD non-treated samples, we found that the gene expression for AC138696.1, macrophage scavenger receptor 1 (MSR1), SMAD family member 6 (SMAD6), CAMP-dependent protein kinase inhibitor beta (PKIB), inka box actin regulator 1 (INKA1), COL23A1, glycoprotein Ib platelet subunit beta (GP1BB), heparan sulfate glucosamine 3-O-sulfotransferase 2 (HS3ST2), CD163, MAF bZIP transcription factor B (MAFB) and Stabilin 1 (STAB1) were significantly upregulated in HC and SCD HU-treated samples but downregulated in non-treated SCD samples. The opposite was found for Fc gamma receptor IIIa (FCGR3A), C2 calcium dependent domain containing 2 (C2CD2), desmocollin 3 (DSC3), histocompatibility minor HB-1 (HMHB1), DIRAS family GTPase 2 (DIRAS2), frizzled class receptor 6 (FZD6), protein tyrosine phosphatase non-receptor type 14 (PTPN14), nuclear receptor subfamily 4 group A member 2 (NR4A2), protein tyrosine phosphatase (PTP), non-receptor type 3 (PTPN3), abelson helper integration site 1 (AHI1), RDH10, Tumor Protein 63 (TP63), GNG4, endosome-lysosome associated apoptosis and autophagy regulator 1 (KIAA1324) and ATPase phospholipid transporting 9A (ATP9A), which were significantly downregulated in HC and SCD HU-treated samples and upregulated in non-treated SCD samples (Figure 2C).

**Figure 1:**
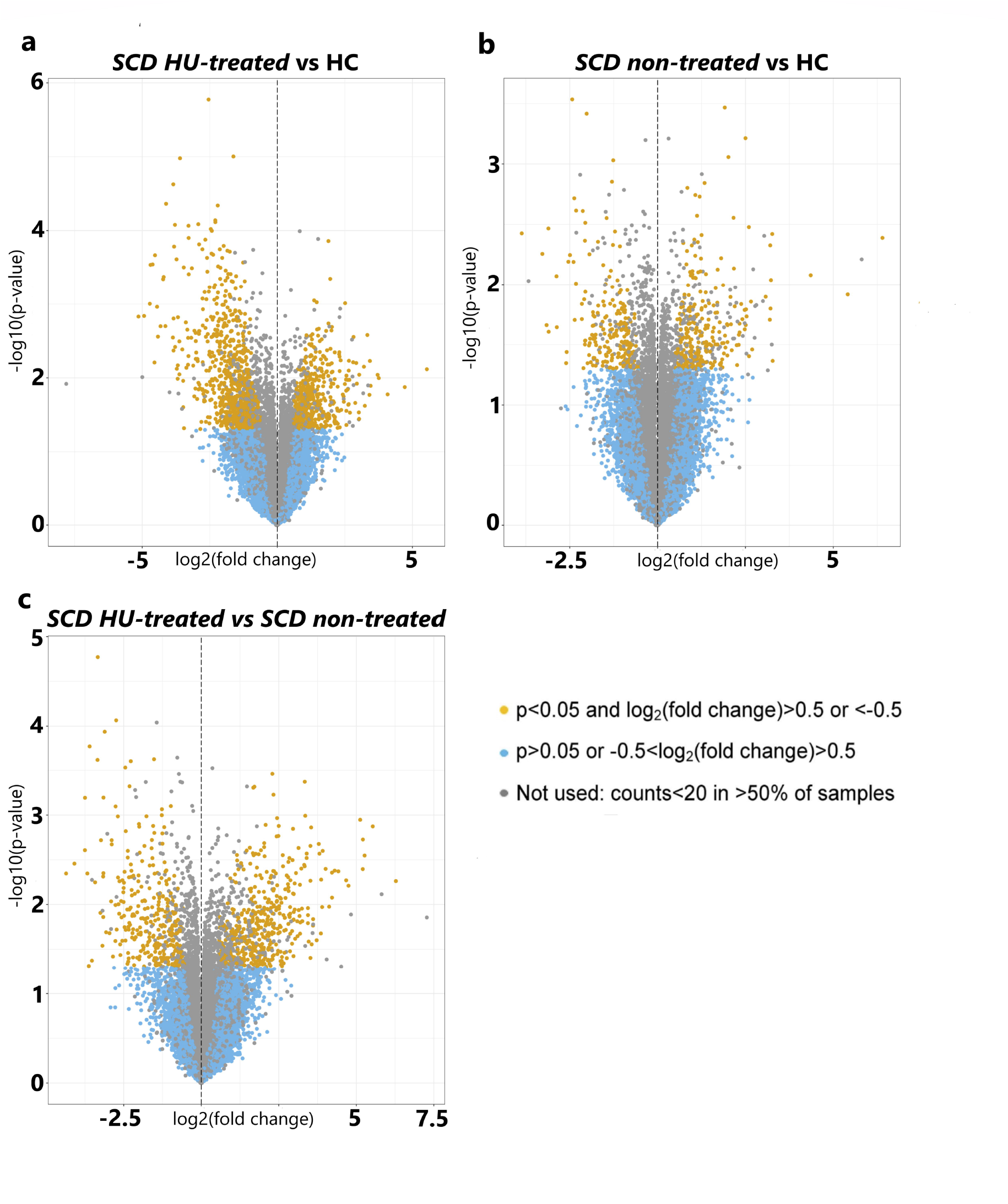
Transcriptomic profiles in SCD patients- and healthy controls (HC) -derived erythroid cells. (a) Volcano plot of 1398 selected DEGs showing up- and downregulated genes in SCD non-treated over HC. (b) Volcano plot of 495 selected DEGs showing up- and downregulated genes in SCD HU-treated over HC. (c) Volcano plot of 786 selected DEGs showing up- and downregulated genes in SCD HU-treated over SCD non-treated.

**Figure 2:**
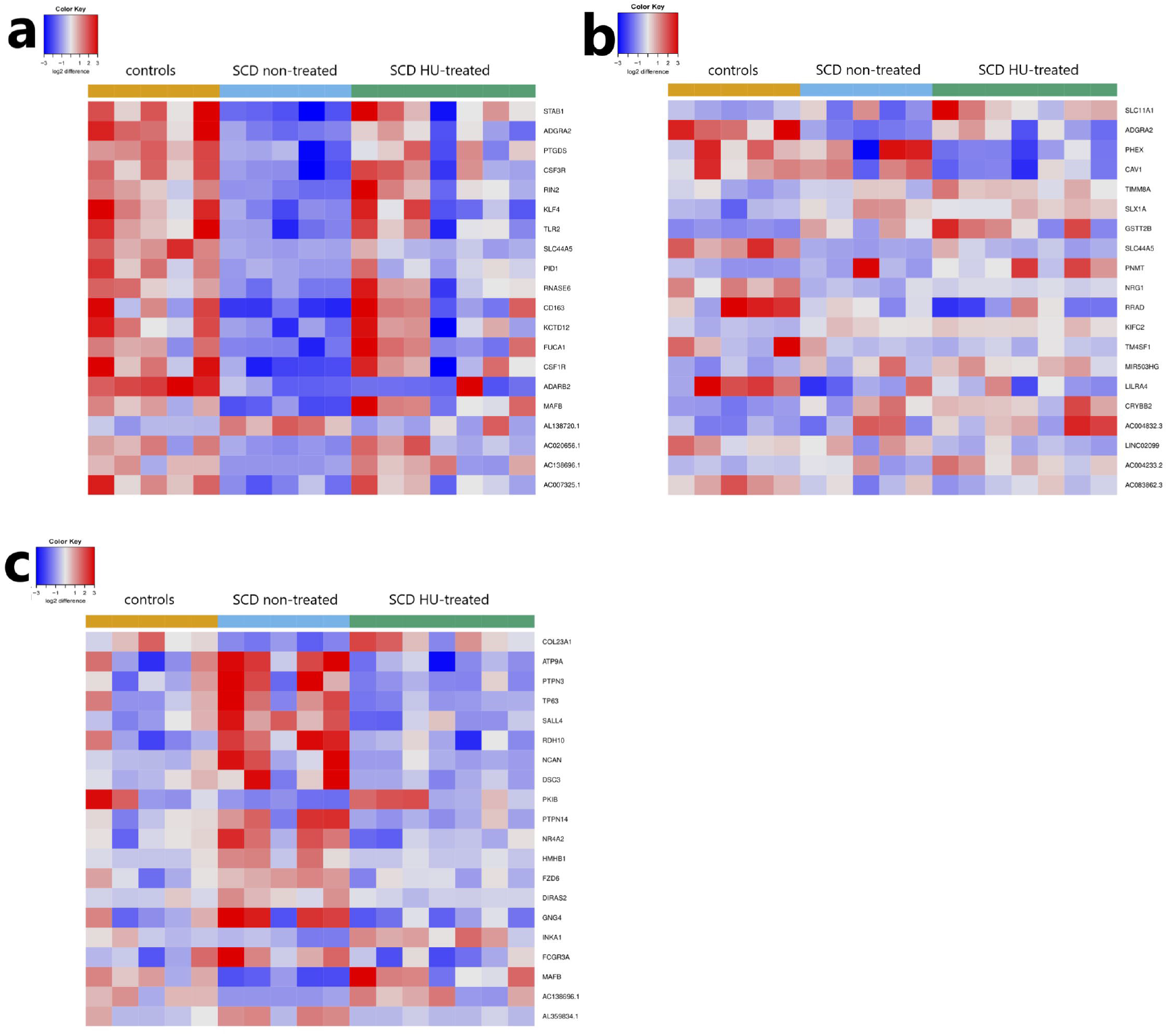
Heatmaps comparing gene expression patterns of the top most significant DEGs in our study groups. (a) SCD non-treated versus healthy controls (b) SCD HU-treated versus healthy controls (c) SCD HU-treated versus SCD non-treated. The colours keys indicate the log2 difference relative to the average expression of all samples within the comparison: in blue are shown downregulated genes and in red are shown upregulated genes. The dendrogram on the left y-axis shows clustering of the genes.

### Molecular pathway analysis

Using the WebGestalt toolkit, we analyzed the RNA sequencing datasets with two approaches^22^: (i) by performing GSEA (Gene Set Enrichment Analysis) in which a list of all detected genes ranked by their average fold change is used to calculate enrichment scores of KEGG (Kyoto Encyclopedia of Genes and Genomes) pathways (significantly enriched pathways are summarized in Figure 3 and Supplementary Figure 3) and (ii) by using DEGs in patient-derived erythroid cells to identify over-represented biological processes and cellular components by ORA (Over-representation Analysis) (Supplementary Figure 2 and Supplementary Table 3). Notably, genes encoding significantly enriched KEGG pathways for Fanconi anaemia and ribosome pathways were upregulated in both SCD non-treated and SCD HU-treated patient-derived erythroid cells in comparison to HC. Both autophagy and mitophagy as well as complement cascades pathways were found downregulated in SCD non-treated group, whereas Toll-like receptor, NF-kappa B signalling were downregulated in SCD HU-treated patient-derived erythroid cells in comparison to HC (Figure 3A-B and Supplementary Figure 3A-B). For HU-treated SCD patient derived erythroid cells in comparison to SCD non-treated, the significantly enriched KEGG pathways included upregulation in complement cascades, phagosome, lysosome and oxidative phosphorylation (Figure 3C, Supplementary Figure 3C). Significantly enriched downregulated pathways included Th1 and Th2 cell differentiation, antigen processing and presentation, T cell receptor signalling pathway and Th1 and Th2 cell differentiation (Figure 3C, Supplementary Figure 3C). Variants in toll-like receptor 2 (TLR2) were found to correlate with the occurrence of bacterial infection in SCD patients.^23^

**Figure 3:**
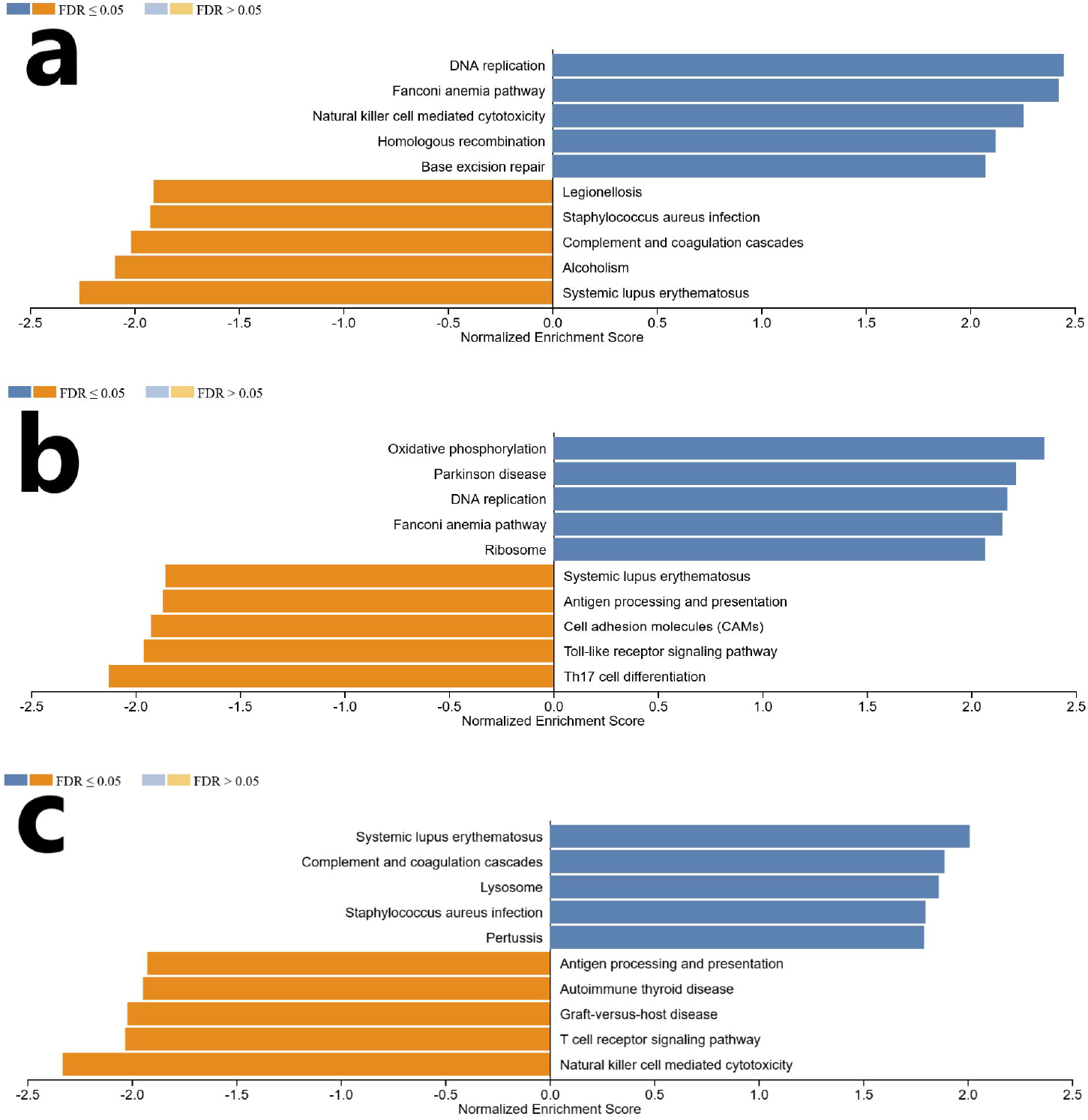
KEGG pathway enrichment analysis depicting significant differentially expressed pathways in our study groups. Each row represents an enrichment function based on FDR (False Discovery Rate). (a) SCD non-treated over HC (b) SCD HU-treated over HC (c) SCD HU-treated over SCD non-treated.

The 10 upregulated and downregulated overlapping DEGs from our 3 study groups are depicted in the Venn Diagram of Supplementary Figure 1 and detailed in Supplementary Table 3. ORA analysis showed the significantly enriched biological processes (e.g., inflammatory response, innate immune response) that are summarized in Supplementary Figure 2 and Supplementary Table 4.

### Selection and validation of top candidate genes

We further selected based on the significance of -log10 (P-value) and the log2 (ratio) values CALN1, KIFC2, MSH5-SAPCD1, GNG4, GSTT2B, RDH10 and FBF1 as our top significant candidate DEGs with biological functions that may contribute to the pathogenesis of SCD (Figure 4). Interestingly, the gene expression of GNG4, MSH5, RDH10 in SCD HU-treated samples were found to be at similar levels as HC suggesting that they may be influenced by the treatment. Whereas the levels of CALN1, KIFC2 and FBF1 in SCD HU-treated and non-treated patients remained at the same levels, significantly different from the levels of HC suggesting that their changed SCD expression may not be affected by the HU treatment (Figure 4).

**Figure 4:**
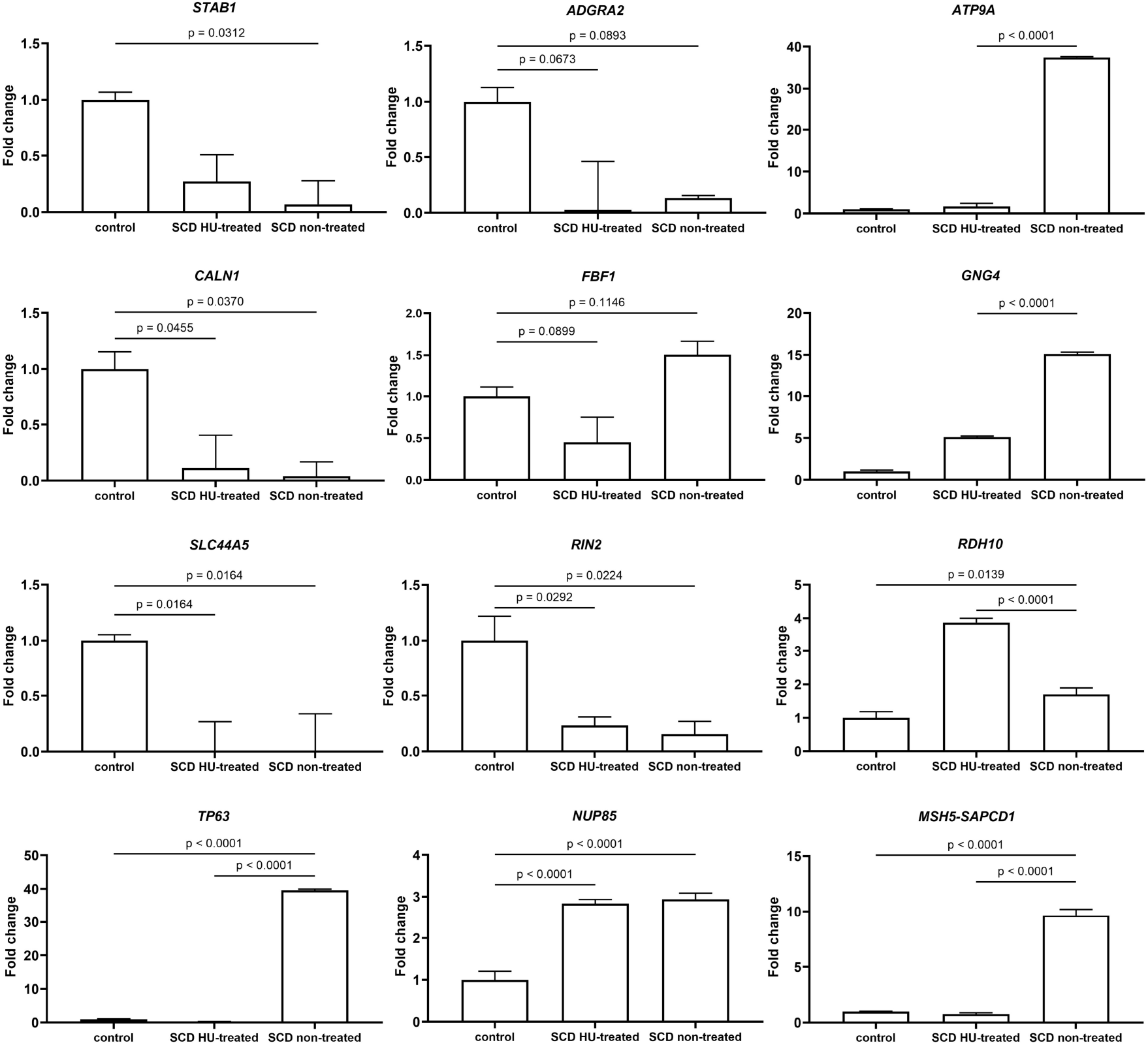
Transcriptomic expression patterns of our top selected genes in our study groups. A significance threshold of 0.05 and a fold change threshold of 0.05 were used for the grouping. The y-axis shows -log10 (P-value) and the x-axis shows log2 (ratio) values.

We then technically validated by qRT-PCR 12 from 17 selected genes (CALN1, NUP85, TP63, Ras and Rab Interactor 2 (RIN2), SLC44A5, PNMT, STAB1, SLX1A, ATP9A, MSH5-SAPCD1, GNG4 and GSTT2B) in 5 SCD non-treated and 5 SCD HU-treated patients. We obtained the same regulation profile in line with the RNA-sequencing data (Figure 5 and Supplementary Table 2).

**Figure 5:**
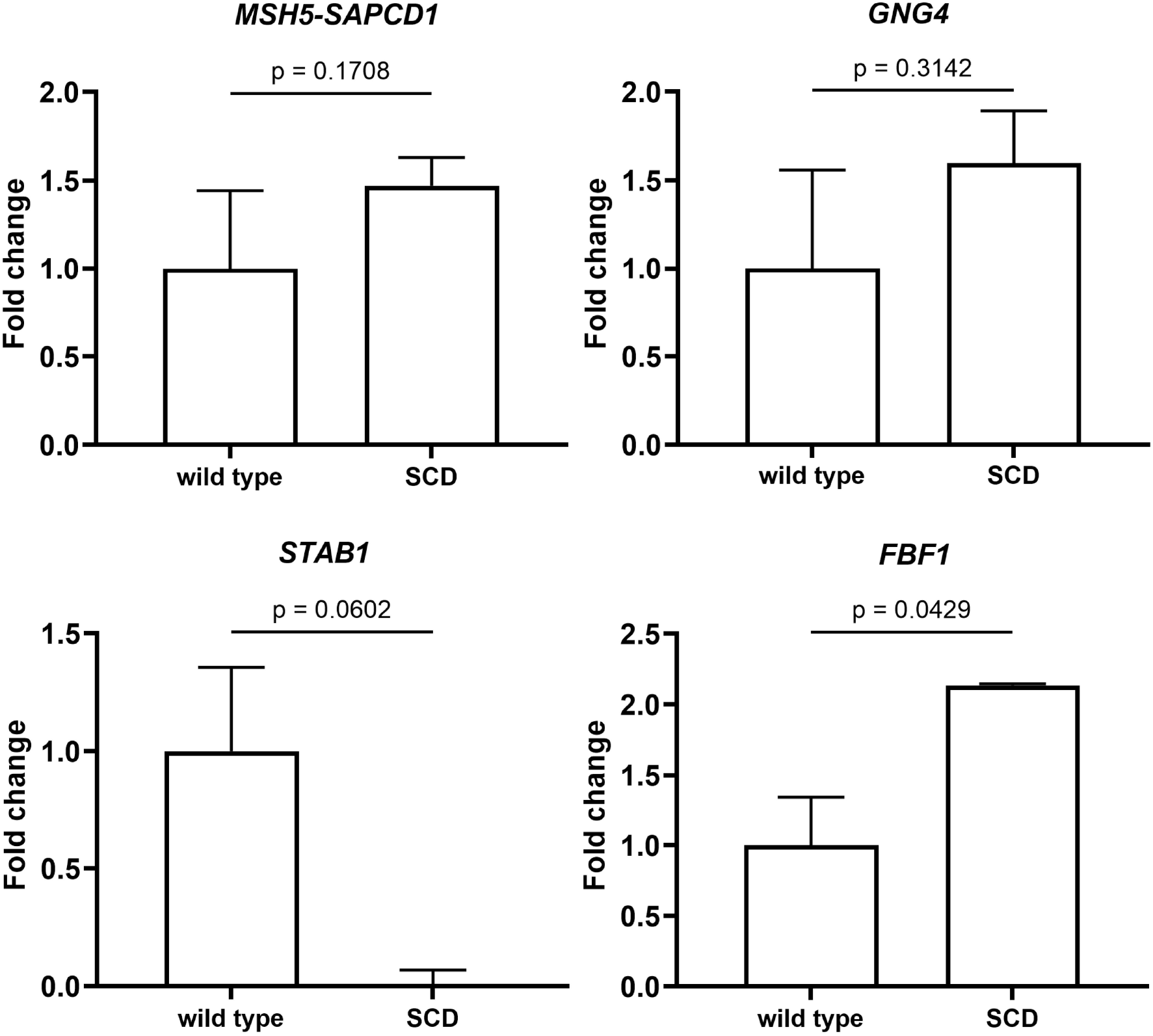
Validation using qRT-PCR of our top selected genes. Total RNA was isolated and subjected to RT-qPCR with 18S rRNA as internal control. Relative mRNA levels of each sample normalized to healthy controls are shown as means (n=4) ± SD. Error bars show SEMs. Statistical analyses were performed using one-way ANOVA followed by multiple comparisons tests to compare the mean ranks between the groups.

We investigated the mRNA levels of these top candidate genes in PBMCs (Supplementary Figure 4). We found a downregulation of mRNA expression of SLC44A5 and CALN1, similar to the erythroid cells from our transcriptomic data, whereas the downregulated mRNA levels of KIFC2 in SCD patients were found the opposite than in RNA sequencing data (Supplementary Figure 4). However, GNG4, NUP85, COL23A1, STAB1, MSH5-SAPCD1, FBF1, RIN2, ATP9A, RDH10 showed no detectable mRNA values in PBMCs. This result suggests the gene patterns specificity of the mRNA gene expression of the erythroid cells in comparison to PBMCs in SCD disease.

### mRNA levels of the top candidate genes in the SCD Berkeley mouse model

To confirm their functional relevance in the SCD disease mechanisms, we investigated the expression of the validated genes in the spleen and bone marrow of the SCD Berkeley mouse model. We observed that the mRNA levels of MSH5-SAPCD, GNG4, FBF1 were upregulated in both spleen and bone marrow of the SCD mouse model compared to the wild type mouse, whereas for STAB1 the mRNA levels were significantly lower (Figure 6 and 7). These results were similar to the mRNA validated expression levels observed in SCD non-treated patients (Figure 5), suggesting that these genes might be interpreted as signatures genes of SCD disease.

**Figure 6:**
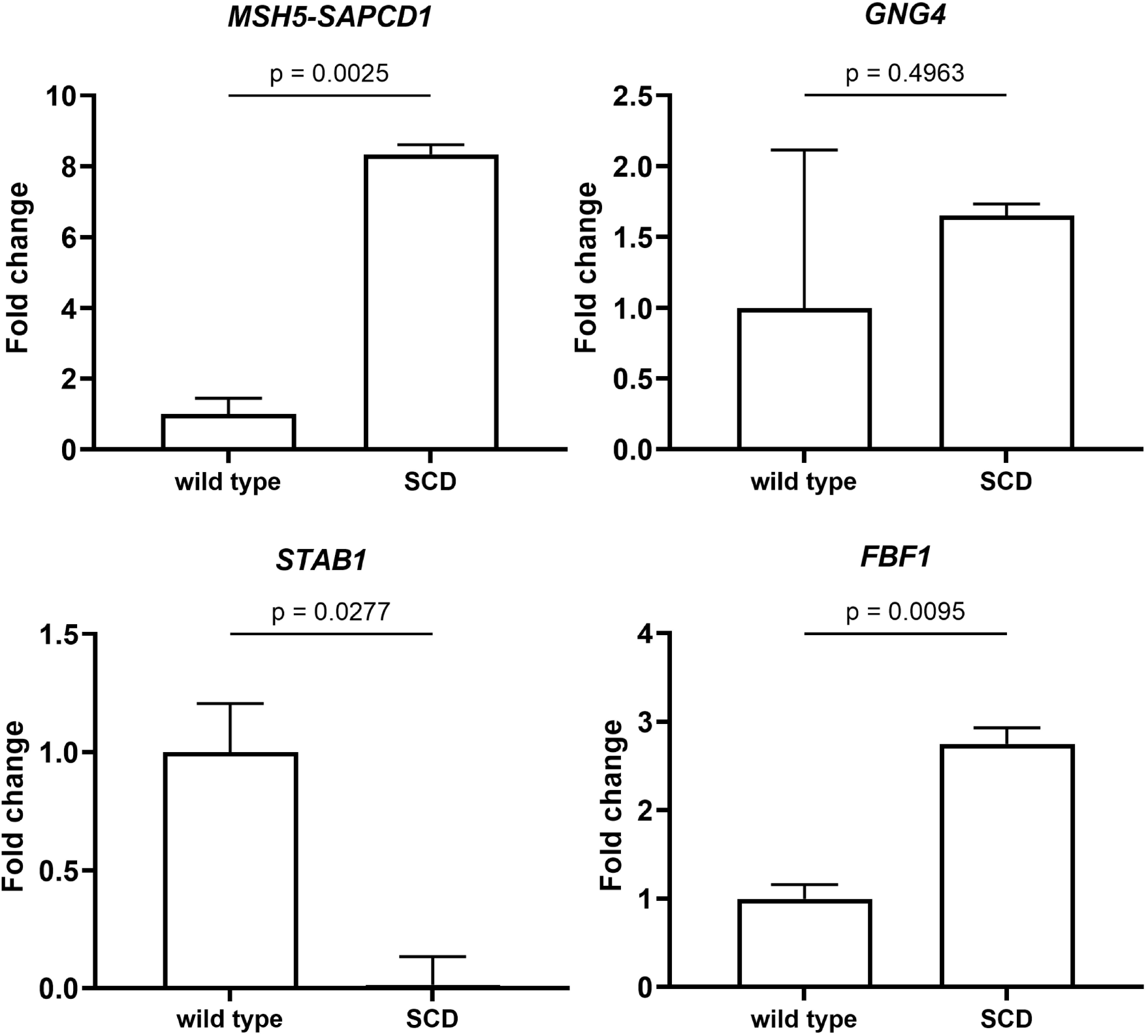
mRNA expression levels of our top selected genes in the bone marrow of the SCD mouse model compared to the wild-type mouse. Total RNA was isolated and subjected to RT-qPCR with 18S rRNA as internal control. Relative mRNA levels of each sample normalized to healthy controls are shown as means (n=4) ± SD. Error bars show SEMs. Statistical analyses were performed using one-way ANOVA followed by multiple comparisons tests to compare the mean ranks between the groups.

**Figure 7:**
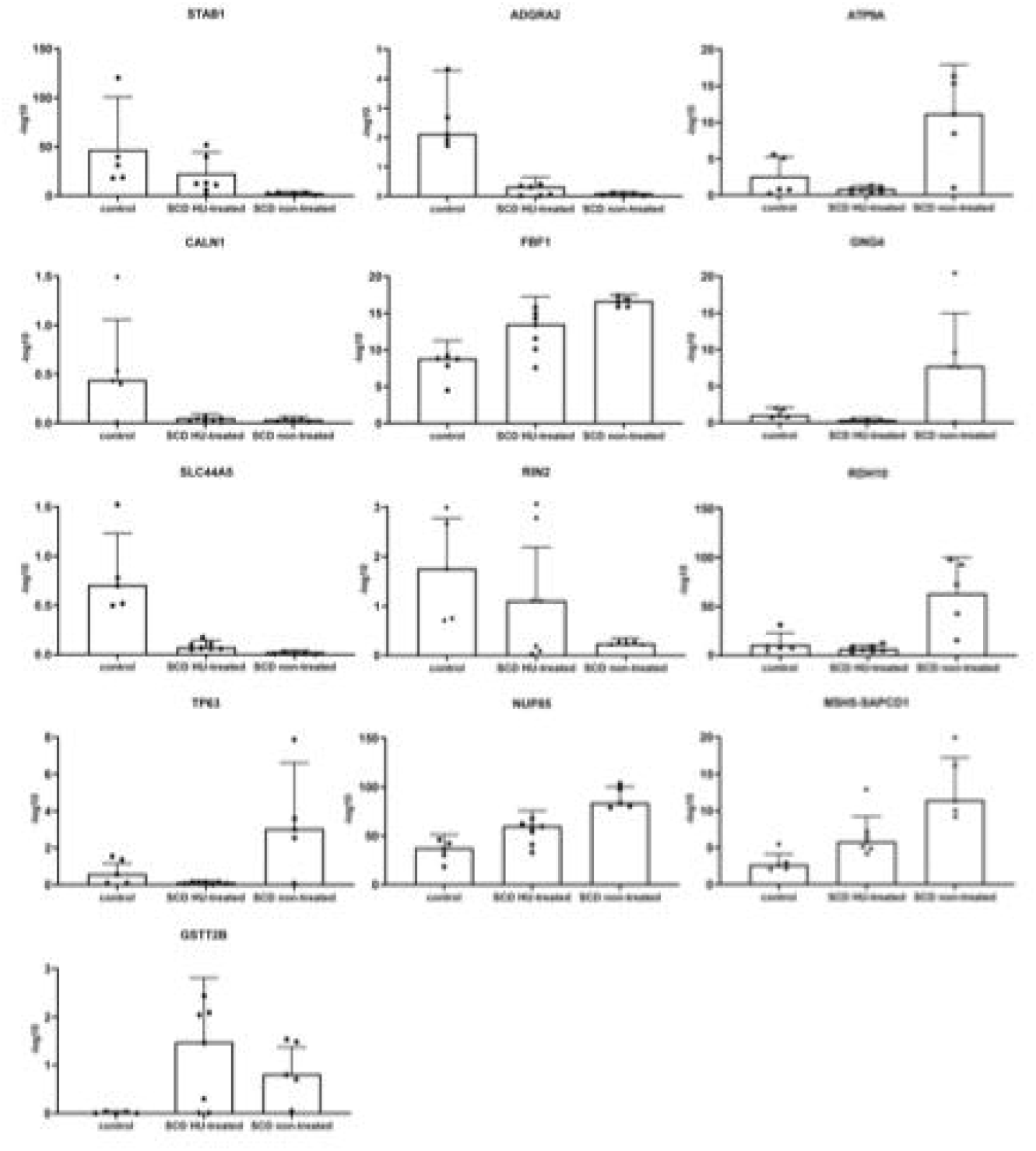
mRNA expression levels of our top selected genes in the spleen of the SCD mouse model compared to the wild-type mouse. Total RNA was isolated and subjected to RT-qPCR with 18S rRNA as internal control. Relative mRNA levels of each sample normalized to healthy controls are shown as means (n=4) ± SD. Statistical analyses were performed using one-way ANOVA followed by multiple comparisons tests to compare the mean ranks between the groups.

## Discussion

We analysed gene expression profiles from children with SCD, treated with HU or untreated, in comparison to HC in an erythropoiesis *ex vivo* model. Overall, we observed significant changes in gene expression and top canonical pathways for SCD, suggesting that our erythroid-specific RNA sequencing might be able to distinguish specific changes according to clinical status and HU treatment effects.

Several previous transcriptomic studies performed in SCD patients revealed prominent roles for proteasome, inflammatory response, innate immunity, haemostasis, haematopoiesis, haemoglobin expression, cytoskeleton, autophagy/mitophagy.^10,24–26^ Two published meta-analyses in whole blood and PBMCs^25,26^ revealed the involvement of innate immunity, response to oxidative stress, erythrocyte development and ubiquitination as most relevant biological processes to SCD. While our study could confirm the contribution by some of these pathways in SCD similar to previous reports (e.g., proteasome, autophagy, immune response, platelet activation), we also generated novel genes and pathways that were not previously associated with the pathogenesis of SCD (NK cytotoxicity, oxidative phosphorylation, Fanconi anaemia pathway, ribosome biogenesis) and might be specific for erythroid cells. ^11^ Moreover, our qRT-PCR analysis in PBMCs cells indicated that there is a gene specificity at the mRNA level of the erythroid cells in comparison to PBMCs in SCD disease, as not all selected genes showed the same patterns. In a more extensive study performed by Gupta et al 2024^1^ it was demonstrated in an in vitro erythropoiesis model that erythroblasts derived from an engineered CRISPR/Cas9 cells could replicated the disease phenotype at the molecular and cellular level. This is in line with our our results confirming that in vitro erythropoiesis may accurately recapitulate similar profiles associated with SCD.

Several studies have shown that SCD patients have a decreased T-lymphocyte count^27^ and found the top significantly enriched Gene Ontology (GO) terms and biological pathways to be associated with T-cell activation and differentiation^26^. T-cell lymphopenia–associated inflammatory responses have been previously linked to the inactivation of PI3Kδ and PI3Kγ^28^, genes that were found to be downregulated in a previous ^26^ meta-analysis in SCD. Concomitant suppression of genes related to adaptive immune responses (B and T cell signalling) was also observed during the two complications of SCD, ACS and VOC in the study from Creary et al.^9^ We found Th1 and Th2 cell differentiation and Th17 cell differentiation to be significantly enriched downregulated KEGG pathways for treated SCD patients in comparison to HC (Figure 3 and Supplementary Figure 3). This result might suggest a contribution to the altered immune response, potentially leading to an increased risk of severe bacterial infections among SCD patients.

A meta-analysis and genome-wide association study by Ben Hamda C. et al.^26^ showed SLC7A5 as one of the top shared upregulated DEGs. In contrast, in our *ex-vivo erythropoiesis* model, SLC44A5 was downregulated and overlapping in SCD non-treated over HC and SCD treated over HC. We can only speculate that these opposite results are specific to the different cellular material that was used for the RNA sequencing in both studies. The solute carrier transporter families are known as metabolic “gatekeepers “ of the cells with an important role in adaptive and innate immunity.^29^ SLC4A1, which belongs to the same solute carrier family, encodes Band 3, an important membrane transporter protein in red cells, with an important role in membrane stability and anione transport in erythropoietic cells.^10^ Whether the regulation of SLC44A5 might play a role in the pathophysiology of SCD, remains to be further elucidated.

Our study showed a downregulation of CALN1 expression in the erythroid cells derived from both SCD treated and non-treated compared to HC, suggesting that HU may not affect CALN1 expression, particularly calcium ion binding. Hänggi et al.^30^ described significantly increased intracellular Ca^2+^ levels in SCD due to an increased number of NMDA receptors causing abnormal Ca^2+^ uptake. Hertz et al.^31^ hypothesized that increased Ca^2+^ levels in RBCs may contribute to an accelerated clearance of RBCs from the blood stream, whereas Wang et al 2021^32^ found a new calcium signalling cascade that is increased in RBCs in SCD patients through the upregulation of lysophosphatidic acid receptor 4. Targeting Ca^2+^ entry mediating molecular pathways with for example Memantine could benefit SCD treatment, as intracellular Ca2+ concentration in human RBCs depends on the abundance and activity of N-methyl D-aspartate receptors (NMDARs) that are present on the RBC ‘s membrane.^33^

GNG4 showed an upregulated gene expression for SCD non-treated patient-derived erythrocytes in comparison to treated patients and HC, suggesting that HU treatment could modify the GNG4 expression. We also found an upregulation of its mRNA expression in the spleen and bone marrow of the SCD mouse model. Previous studies showed that after downregulating the expression of GNG4 *in vitro*, proliferation, migration and invasion of colon cancer cells were significantly reduced and the cell cycle was blocked in the S phase.^34^ Interestingly, these results may suggest a common benefit in inhibiting the GNG4 expression, as seen in our DEG data from treated patients.

FBF1 showed an upregulated gene expression for both SCD treated and non-treated patient-derived erythroid cells in comparison to HC, in SCD HU-treated being downregulated compared to SCD non-treated, suggesting that HU may affect FBF1 expression. We also found an upregulated mRNA expression in the spleen and bone marrow of the SCD mouse model. Fas and its ligand (FasL) interactions are known as major inducer of apoptosis and of inflammatory response.^35^ In line with our results, high levels of sFas/sFasL ratio were previously reported in SCD, this ratio being used as a marker for vascular dysfunction and assessing renal dysfunction in SCD patients, in relation to inflammation, iron deposition and albuminuria.^35^

Our results indicated that the expression of STAB1 was downregulated not only in SCD HU-treated and non-treated groups compared to HC, but as well as in the spleen and bone marrow of the SCD mouse model. STAB1 is expressed in alternatively activated macrophages, with an important role in tissue homeostasis and prevention of autoimmunity.^36,37^ As blocking STAB1 in macrophages results in defective engulfment of aged red blood cells,^37^ we could assume that STAB1 could play a role in their clearance in SCD. Future studies are required to clarify this issue.

GSTT2B showed an upregulated gene expression for SCD treated patient-derived erythroid cells in comparison to HC and non-treated patients, suggesting a possible influence of HU on GSTT2B expression, as HU can improve antioxidant defences of SCD patients.^38^ GSTT2B is a member of a superfamily of proteins that catalyze the conjugation of reduced glutathione to a variety of electrophilic and hydrophobic compounds.^39^ A previous study indicated that reduced glutathione in red cells is an important antioxidant whose depletion together with the low reduced glutathione levels in HbS cells may contribute to the pathophysiology of SCD,^40^ by not yet known mechanisms.

The mechanisms of the induction of HbF induction by HU treatment and its effects on the global transcriptome are not yet fully elucidated.^41^ Chondrou V. et al. (2017)^41^ performed a whole transcriptome analysis in erythroid cells derived from human hematopoietic tissues of various developmental stages where they identified VEGFA gene to be associated with elevated fetal haemoglobin levels in β-type hemoglobinopathy patients. They indicated that VEGFA genomic variants were associated with disease severity in β-thalassemia patients and HU treatment efficacy in SCD/β-thalassemia compound heterozygous patients. In line with this, we also found a significant downregulation of the mRNA levels of RIN2 in our SCD treated and non-treated groups compared to HC, an activator of early endosomal GTPase Rab5C which stabilizes the VEGFR2 levels, crucial for VEGF signalling through ERK and PI3-K pathways.^42^ In a global transcriptomic analysis of mouse embryonic stem cells in response to HU-treatment, Cui P. et al. (2010)^43^ found a suppressed transcriptional activity after HU-treatment exposition. HU-treatment altered multiple key cellular pathways, including cell cycle, apoptosis, inflammatory response, DNAs- and alternative splicing mechanisms and suppressed non-coding RNA expression. Similar to the upregulation of DNA repair and replication pathways observed in our analysis, many DNA damage repair enzymes were upregulated in HU-treated embryonic stem cells.^48^

There is scarce data available regarding the transcriptomic gene profile and gene regulation mechanisms in SCD transgenic mouse models in comparison to human SCD patients. Previous work in sickle transgenic mouse indicated the role of NF-kB signalling in blood mononuclear cells in regulating endothelial tissue factor expression^44^ or that the loss of NRF2, as a regulator of oxidative stress and heme activator, could exacerbate the pathophysiology of SCD disease.^45^ We observed that some of the genes that demonstrated an upregulation of the mRNA levels in the human SCD erythroid cells were also upregulated in the bone marrow and spleen from the SCD mouse model (e.g., STAB1, GNG4, FBF1). These results suggest that SCD disease might share some common molecular mechanisms in both human cells and mouse model.

Taken together, we could identify a new subset of genes associated for the first time with SCD. These findings may encourage future characterization of their roles in SCD pathogenesis and could serve to identify new treatment possibilities in SCD patients and other haemoglobinopathies.

## Materials and methods

### Patients recruiting and ethical considerations

All patients were recruited at the Children ‘s Hospital Zurich, Switzerland. All experimental procedures were approved by the local ethical committee (Cantonal Ethics Committee Zurich, Switzerland, https://www.zh.ch/de/gesundheitsdirektion/ethikkommission.html). All healthy blood donors as well as SCD patients, or their legal guardians gave written informed consent prior to their enrolment in this study. All methods were carried out in accordance with the appropriate guidelines and regulations. As patients were under the legal age of consent, informed consent was obtained from parent/legal guardians prior to the start of this study. Details of patient enrollment and sample collection, have been previously published.^10^ All methods are reported in accordance with ARRIVE guidelines.

### Erythroblast expansion and differentiation (ex-vivo erythropoiesis)

PBMCs were isolated from heparin blood using the SepMateTM PBMC isolation tubes from STEMCELL Technologies Inc. according to manufacturer ‘s protocol. PMBCs layer was removed into a new Falcon tube, diluted 1:1 with RPMI 1640 medium (Thermo Fischer Scientific) and centrifuged at 1000g for 10 minutes. After removal of the supernatant, resuspension of the pellet and repeat centrifugation, the pellet of PBMCs was divided, one part was freezed at -80°C for further RNA extraction, the remaining part was then cultured in a two-phase liquid system as previously described by our group.^11,46^

### Transgenic and control mice

The Berkeley mouse model (*JAX stock #003342*) homozygous for the alpha-globin null allele, homozygous for the beta-globin null allele and carrying the sickle transgene (Hba^0/0^ Hbb^0/0^ Tg(HuminiLCRα1^G^γ^A^γδβ^S^) was pursued from Jackson Laboratory (Strain 003342) and was kindly provided to us by Dr. Dominik Schaer and Dr. Florence Vallelian, University Hospital Zurich (n=4). All breeding colonies were housed and bred in the specific pathogen-free (SPF) animal facility at the Laboratory Animal Services Center (LASC) of the University of Zurich in individually ventilated cages. Mice were housed under a 12/12-h light/dark cycle in accordance with international guidelines. Mice that were 10-12 weeks old were used for all experiments. Only healthy, well-conditioned mice with uncompromised and groomed fur were included as experimental animals. All experimental protocols were reviewed and approved by the Veterinary Office of the Canton of Zurich (ZH120 2021). All animal experiments were performed according to Swiss and ARRIVE guidelines. The animals were dissected to expose the spleen and femur bone marrow. The tissues were mechanically processed for RNA extraction.

### RNA-sequencing Data Analysis

Total RNA was extracted from cultured erythroid cells and PBMCs using RNeasy mini kit from Qiagen, Germany, as described by the manufacturer and previously.^17^

The libraries were prepared with Illumina ‘s TruSeq Stranded mRNA kit using 300ng input total RNA. RNA sequencing was performed on an Illumina NovaSeq 6000 instrument at the Functional Genomics Center Zurich. The raw reads were cleaned by removing adapter sequences, trimming low quality ends, and filtering reads with low quality (phred quality <20) using Fastp (Version 0.20).^47^ Pseudo sequence alignment of the resulting high-quality reads to the human transcriptome (genome build GRCh38.p10, gene annotations based on GENCODE release 32) and quantification of transcript expression was carried out using Kallisto (Version 0.46.1)^48^.To detect DEGs, we applied a count based negative binomial model implemented in the software package EdgeR (Rversion: 4.0.3, EdgeR version: 3.32).^49^ The differential expression was assessed using the GLM framework and applying a quasi-likelihood F-test. The resulting p-values were adjusted with the Benjamini and Hochberg method.

### RT-qPCR Analysis

The RNA was reverse transcribed using the QuantiTect Reverse Transcription Kit (Qiagen) as recommended by the manufacturer. RT-qPCR was performed using PowerUp SYBR Green Master Mix (Thermofisher Scientific, Switzerland, cat. A25742) and primers with specific sequences as mentioned in Supplementary Table 1.

### Statistical Analysis

Our data was statistically analyzed by using Prism 9.5.0 (GraphPad, Software CA, USA). Data was analysed using t-Tests or One-way ANOVA, followed by multiple comparisons tests to compare the means between the groups and p value < 0.05 was considered significant.

## Acknowledgements

We would like to thank to Stacey Shllaku, Nadja Schulthess and Alessandra Bosch for the technical support.

## Contributions

The study was conceived and coordinated by FDF and MS. The experiments were designed by MS, DJS, FV and FDF. The experiments were conducted by MT and FDF. Bioinformatic analysis was conducted by LO, MT and FDF. Data analysis was conducted by MT and FDF. The project was supervised by FDF and MS. The manuscript was drafted and refined by MT and FDF with final revisions from MS, LO, DJS and FV.

## Data availability

The RNA-seq data are available under ENA repository under the accession ID: PRJEB69480.

## Funding

This study was supported from research grants from University of Zurich (MS) and Jacques & Gloria Gossweiler Foundation grant (FDF and MS).

## Competing interests

The authors declare no competing interests.

## Notes

### Competing Interest Statement

The authors have declared no competing interest.

### Summary of Updates

Figures updated and text has been improved

