## Supplementary Data for "Transcriptomic gene profiles in an *ex vivo* model of erythropoiesis to unravel molecular pathomechanisms of Sickle Cell Disease"

**Supplementary Table 1.** Primers used for RT-qPCR.

| Gene Name | Species | Primer Sequence |
| --- | --- | --- |
| CALN1_forward | human | 5’-CCG CAC ACA GAG TCT CAG -3’ |
| CALN1_reverse | human | 5’-CAT GGT GGA ACG GCA TCT -3’ |
| NUP85_forward | human | 5’-GCA CAT TGA GCG GAT ACC T -3’ |
| NUP85_reverse | human | 5’-CCC TGA GGA ACC TGT CTG A -3’ |
| KIFC2_forward | human | 5’-AGC CAG GAG GAG GTC TTC -3’ |
| KIFC2_reverse | human | 5’-CCT GAC AGC CTC ATT GTA GAT -3’ |
| MSH5-SAPCD_forward | human | 5’-TGG CAA GGA GGT CTC AGA -3’ |
| MSH5-SAPCD_reverse | human | 5’-AGG ACA CTG GAA GGA CTC T -3’ |
| GNG4_forward | human | 5’-TCA CCT CTC ATC TGA CGA CTG -3’ |
| GNG4_reverse | human | 5’-TGC CTG GGA GAC CTT GAC -3’ |
| GSTT2B_forward | human | 5’-CTG TGG GTC CAG GTG TTG -3’ |
| GSTT2B _reverse | human | 5’-CAG TTC ATA GCC GAG AGC C-3’ |
| RDH10_forward | human | 5’-AGT CTC AGT CCT GGT CAA TAA TG-3’ |
| RDH10_reverse | human | 5’-TGG CAC AGT AAT CCT CAA CTC -3’ |
| FBF1_forward | human | 5’-CAC ATA CCA GAG ACA CCA CAG -3’ |
| FBF1_reverse | human | 5’-CCG TCC AGG TCC TTC ATG -3’ |
| SLX1A_forward | human | 5’-CCT GCT CTA CTG CCT GAA C -3’ |
| SLX1A_reverse | human | 5’-AGC CCA CTC AAA CCG AAG -3’ |
| RIN2_forward | human | 5’-GAG AAG GAG GCT ATT ACT TGA CAA -3’ |
| RIN2_reverse | human | 5’-CAA CTC GGA GGT AAT TCT GGA A -3’ |
| COL23A1_forward | human | 5’-GCG TCT GAC AGC CTA CAG -3’ |
| COL23A1_reverse | human | 5’-GTC TGC CCT TCT CTC CTT TG -3’ |
| STAB1_forward | human | 5’-GGA CAA CAT GAC GCT GAG T -3’ |
| STAB1_reverse | human | 5’-ACA ATG ATA CGG CTA ACC ACA A -3’ |
| PNMT_forward | human | 5’- CCT ACC TCC GCA ACA ACT -3’ |
| PNMT_reverse | human | 5’-AAT CTG TCA TGG TGA TGT CCT C -3’ |
| ADGRA2_forward | human | 5’- CCG TTA CCC TGC TCT TGA G -3’ |
| ADGRA2_reverse | human | 5’-GAG GTG AGA CAG CCA ATC C -3’ |
| ATP9A_forward | human | 5’- GAT CCT GAA CTT CAC CAT CCT AC -3’ |
| ATP9A_reverse | human | 5’-TGC CAC ACT CTT CCT CCA -3’ |
| 18S_forward | human | 5’-GTT CCG ACC ATA AAC GAT GCC -3’ |
| 18S_reverse | human | 5’-TGG TGG TGC CCT TCC GTC AAT -3’ |
| NUP85 | human | BIO-RAD (PrimePCR SYBR Green Assay)  (UniqueAssayID: qHsaCID0018193) |
| SLC44A5 | human | BIO-RAD (PrimePCR SYBR Green Assay)  (UniqueAssayID: qHsaCED0057217) |
| TP63 | human | BIO-RAD (PrimePCR SYBR Green Assay)  (UniqueAssayID: qHsaCID0036332) |
| CALN1 | human | BIO-RAD (PrimePCR SYBR Green Assay)  (UniqueAssayID: qHsaCED0047583) |
| GNG4 | human | BIO-RAD (PrimePCR SYBR Green Assay)  (UniqueAssayID: qHsaCED003675) |
| GSTT2B | human | BIO-RAD (PrimePCR SYBR Green Assay)  (UniqueAssayID: qHsaCID0038698) |
| GNG4 | mouse | BIO-RAD (PrimePCR SYBR Green Assay)  (UniqueAssayID: qMmuCID0018380) |
| MSH5 | mouse | BIO-RAD (PrimePCR SYBR Green Assay)  (UniqueAssayID: qMmuCID0022924) |
| KIFC2 | mouse | BIO-RAD (PrimePCR SYBR Green Assay)  (UniqueAssayID: qMmuCID0022929) |
| FBF1 | mouse | BIO-RAD (PrimePCR SYBR Green Assay)  (UniqueAssayID: qMmuCID0021443) |
| 18S_forward | mouse | 5’- CTT AGA GGG ACA ATC CAA -3’ |
| 18S_reverse | mouse | 5’- ACG CTG AGC CAG TCA GTG TA -3’ |

**Supplementary Table 2.** Summarized qRT-PCR validation results of top significant genes from our transcriptomic study.

| Gene Symbol and Name | Transcriptomic Results | Validation Results |
| --- | --- | --- |
| CALN1  (calneuron 1) | Downregulated in SCD HU-treated and non-treated vs HC | validated |
| NUP85  (nucleoporin 85) | Upregulated in SCD HU-treated and non-treated vs HC | validated |
| TP63  (tumor protein 63) | Downregulated in SCD HU-treated vs SCD non-treated | validated |
| RIN2  (Ras and Rab interactor 2) | Downregulated in SCD non-treated vs HC | validated |
| KIFC2  (kinesin family member C2) | Upregulated in SCD HU-treated and non-treated vs HC | not validated |
| MSH5-SAPCD1  (MSH5-SAPCD1 readthrough (NMD candidate) | Upregulated in SCD non-treated vs HC | Validated, significant in mouse spleen, statistically significant in mouse bone marrow |
| COL23A1  (collagen type XXIII alpha 1 chain) | Downregulated in SCD non-treated vs HC | not validated |
| SLC44A5  (solute carrier family 44 member 5) | Downregulated in SCD HU-treated and non-treated vs HC | validated |
| GNG4  (G protein subunit gamma 4) | Downregulated in SCD HU-treated vs SCD non-treated | Validated, not statistically significant in mouse spleen and in mouse bone marrow |
| GSTT2B  (glutathione S-transferase theta 2B) | Upregulated in SCD HU-treated vs HC | validated |
| PNMT  (phenylethanolamine N-methyltransferase) | Upregulated in SCD HU-treated vs HC | validated |
| STAB1  (stabilin 1/ clever-1) | Downregulated in SCD non-treated vs HC | Validated, significant in mouse spleen, statistically significant in mouse bone marrow |
| SLX1A  (SLX1 homolog A, structure-specific endonuclease subunit) | Upregulated in SCD HU-treated and non-treated vs HC | validated |
| ADGRA2  (adhesion G protein-coupled receptor A2) | Downregulated in SCD HU-treated and non-treated vs HC | Not validated |
| ATP9A  (ATPase phospholipid transporting 9A (putative)) | Downregulated in SCD HU-treated vs SCD non-treated | validated |
| RDH10  (retinol dehydrogenase 10) | Downregulated in SCD HU-treated vs SCD non-treated | Not validated |
| FBF1  (Fas binding factor 1) | Upregulated in SCD HU-treated and non-treated vs HC | Not validated, significant in mouse spleen, statistically significant in mouse bone marrow |

**Supplementary Table 3.** Overlapping up- and downregulated DEGs from Supplementary Figure 1 (upregulated genes are red and downregulated genes are blue).

| Overlapping up- and downregulated DEGs | | | |
| --- | --- | --- | --- |
| Gene Symbol | Trend | Gene Symbol | Trend |
| ADGRA2 | DOWNREGULATED | NUP85 | UPREGULATED |
| FCN1 | DOWNREGULATED | SLX1A | UPREGULATED |
| ONECUT2 | DOWNREGULATED | SLFN13 | UPREGULATED |
| STEAP4 | DOWNREGULATED | FBF1 | UPREGULATED |
| CDKN1C | DOWNREGULATED | CRYBB2 | UPREGULATED |
| SLC44A5 | DOWNREGULATED | MSH5-SAPCD1 | UPREGULATED |
| GPM6A | DOWNREGULATED | CCER2 | UPREGULATED |
| NRG1 | DOWNREGULATED | AL035461.2 | UPREGULATED |
| RRAD | DOWNREGULATED | F8A3 | UPREGULATED |
| CALN1 | DOWNREGULATED | CU638689.4 | UPREGULATED |

**Supplementary Table 4.** Over-represented gene ontology terms (ORA) among DEGs in SCD non-treated vs HC, SCD HU-treated vs HC, SCD versus HC in erythroid cells.

| **Gene Ontology** | **Description** | **Gene Ratio** | **p-value** |
| --- | --- | --- | --- |
| **SCD_nonTreated–over–HC: Biological Process** | | | |
| <GO:0006281> | DNA repair | 12/75 | 2.063409e-06 |
| <GO:0032508> | DNA duplex unwinding | 5/75 | 9.247631e-05 |
| <GO:0006974> | cellular response to DNA damage stimulus | 11/75 | 1.509270e-04 |
| <GO:0006260> | DNA replication | 6/75 | 3.747499e-04 |
| <GO:0006954> | inflammatory response | 36/257 | 8.904680e-19 |
| <GO:0043312> | neutrophil degranulation | 42/257 | 6.514479e-17 |
| <GO:0045087> \| | innate immune response | 31/257 | 5.391094e-11 |
| **SCD_Treated-over-HC: Biological Process** | | | |
| <GO:0002250> | adaptive immune response | 6/24 | 1.733475e-05 |
| <GO:0006958> | complement activation, classical pathway | 4/24 | 2.751195e-05 |
| <GO:0045087> | innate immune response | 6/24 | 6.819515e-05 |
| **SCD_Treated–over–SCD_nonTreated: Biological Process** | | | |
| <GO:0006954> | inflammatory response | 14/96 | 2.938618e-08 |
| <GO:0006952> | defense response | 8/96 | 7.415814e-08 |
| <GO:0001942> | hair follicle development | 4/74 | 1.400842e-05 |
| <GO:0050776> | regulation of immune response | 7/74 | 5.472021e-05 |
| **SCD-over-HC: Biological Process** | | | |
| <GO:0006281> | DNA repair | 7/43 | 0.0002678932 |
| <GO:0006974> | cellular response to DNA damage stimulus | 7/43 | 0.0013140272 |
| <GO:0032508> | DNA duplex unwinding | 3/43 | 0.0023127094 |
| <GO:0006310> | DNA recombination | 3/43 | 0.0042224542 |

**Supplementary Figure 1.** **Venn Diagram showing number of overlapping and non-overlapping DEGs**. *p* <0.01 for SCD non-treated vs HC and SCD HU-treated vs HC (upregulated genes are in red and downregulated genes are in blue).

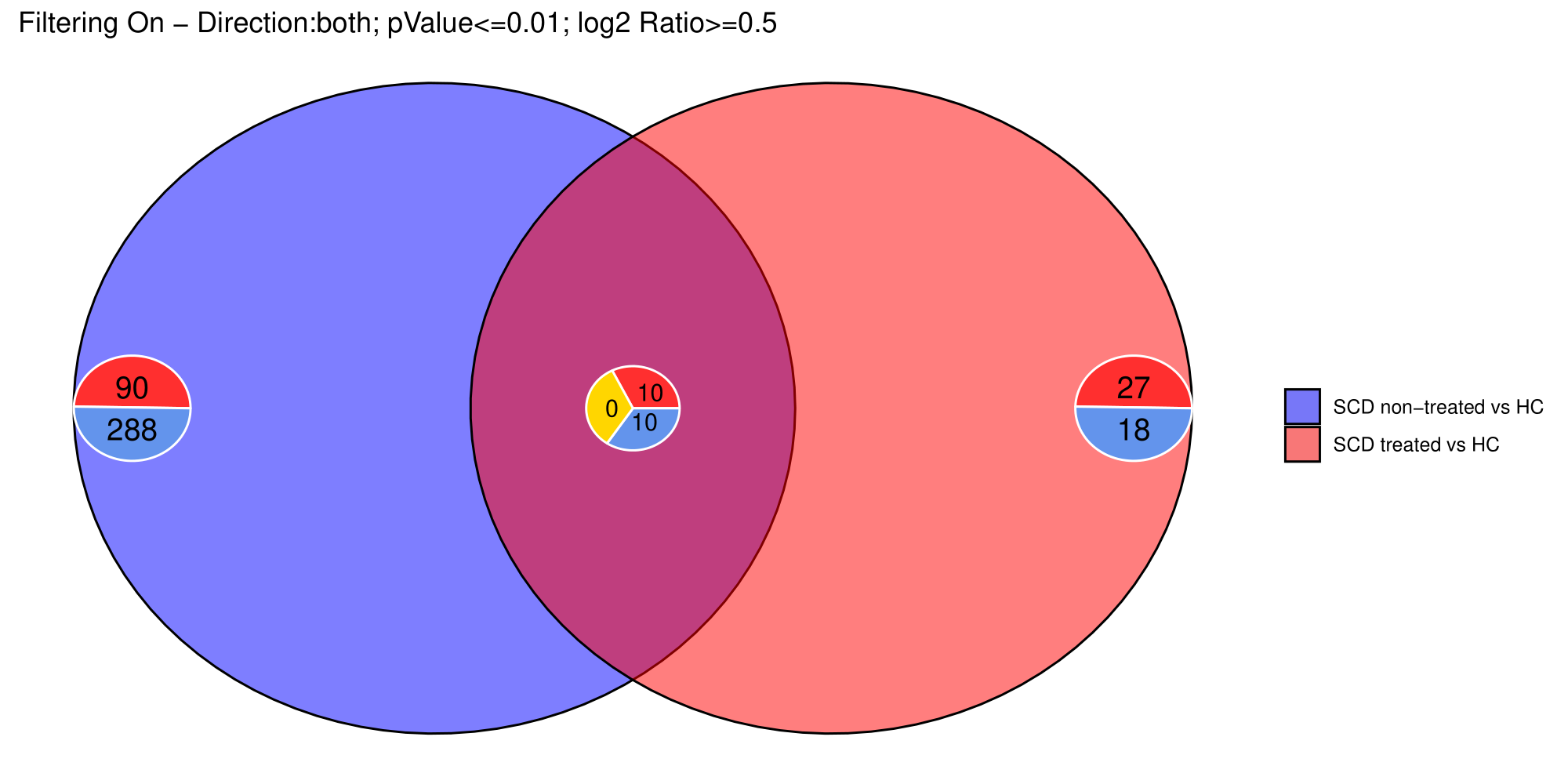

**Supplementary Figure 2.** **Overrepresentation analysis (ORA) showing top pathways enriched by the genes of the chemokine-cytokine, scavanger receptor, extracellular matrix activity.** Each pathway is indicated according to their p value by a red (most significant p values) or orange circle accordingly. The circle size corresponds to the number of genes enriching that pathway. The results are shown for HU-treated vs non-treated SCD patient groups taken from our transcriptomic profiling, including top DEGs COL23A1 or STAB1.

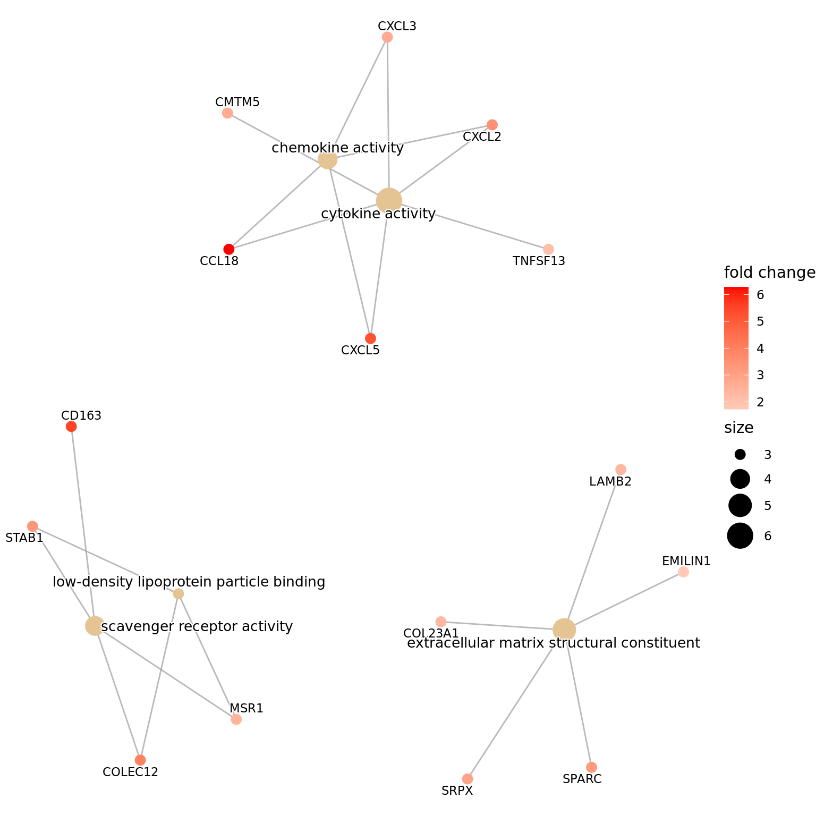

**Supplementary Figure 3.** **Bar chart of KEGG pathway enrichment analysis depicting top 10 significant differentially expressed pathways in our study groups.** Each row represents an enrichment function based on FDR. A) SCD non-treated over HC. B) SCD HU-treated over HC. C) SCD HU-treated over SCD non-treated.

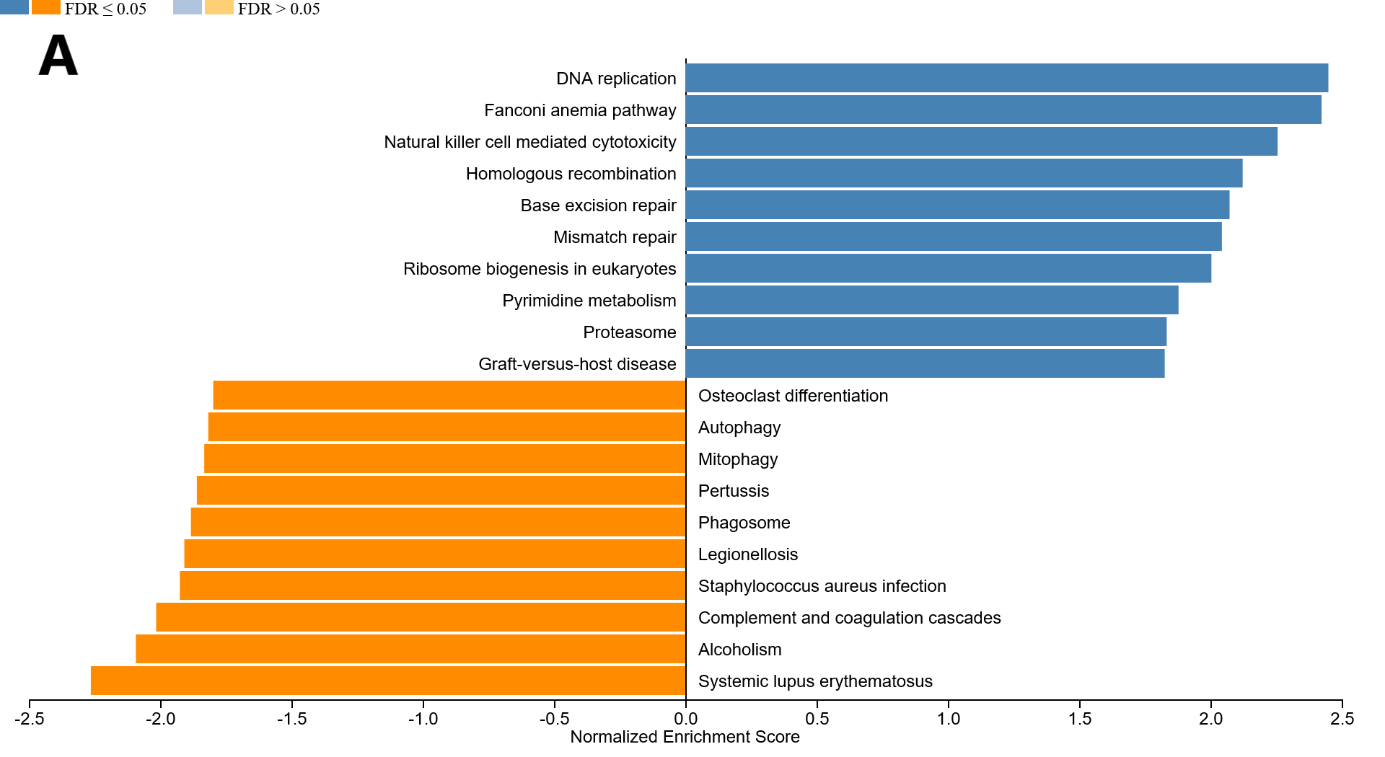

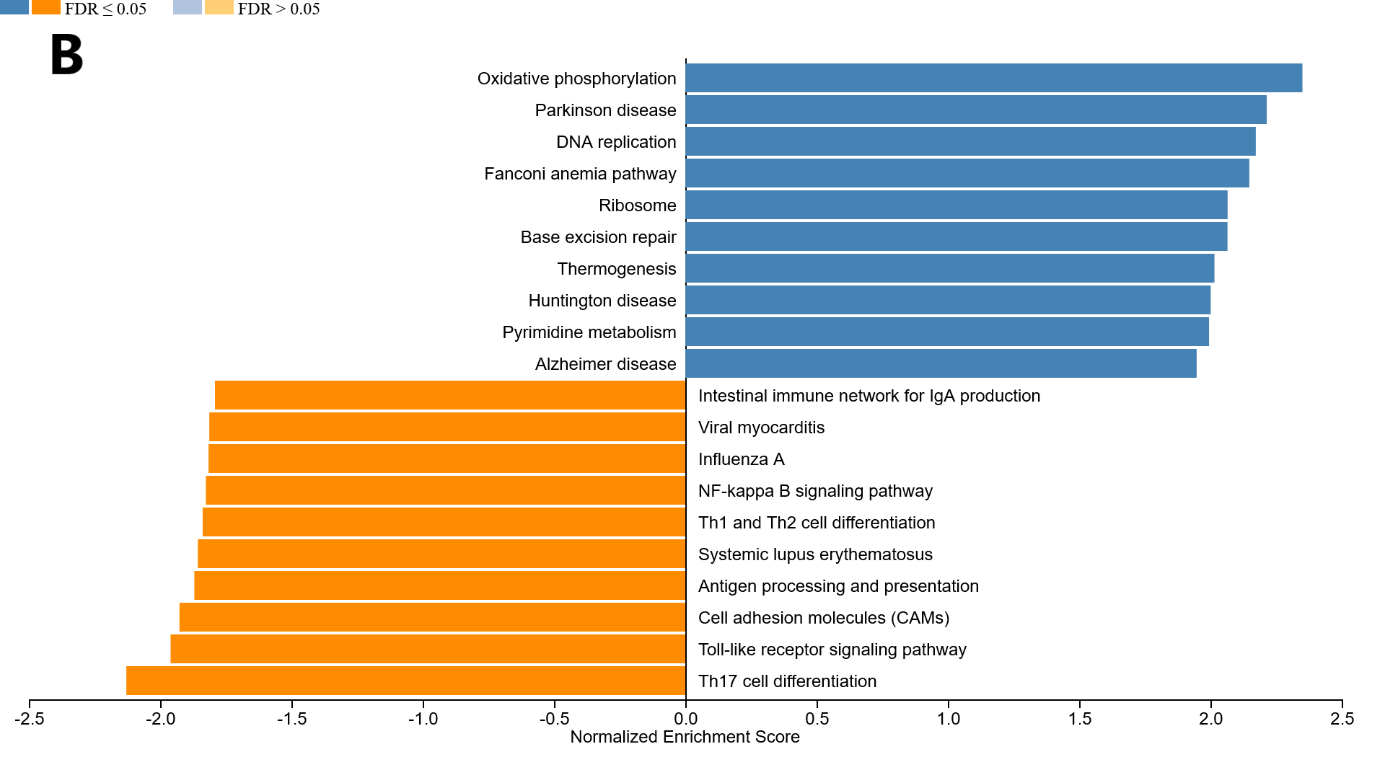

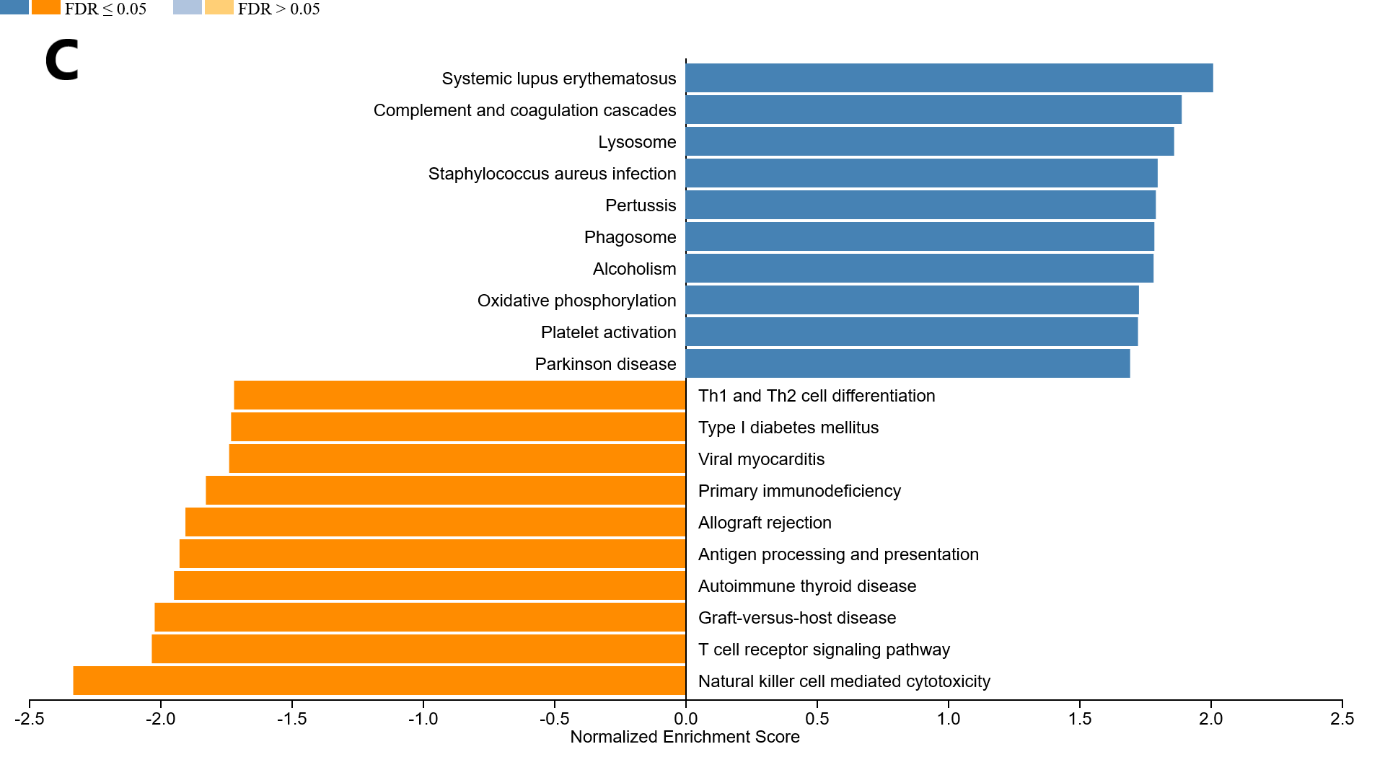

**Supplementary Figure 4. mRNA expressions for SLC44A5, CALN1 and KIFC2 in PBMCs.** Gene expression was measured in four independent replicates per subject by qRT-PCR. t-tests were performed for statistical analysis. Data is shown as mean ± SEM.

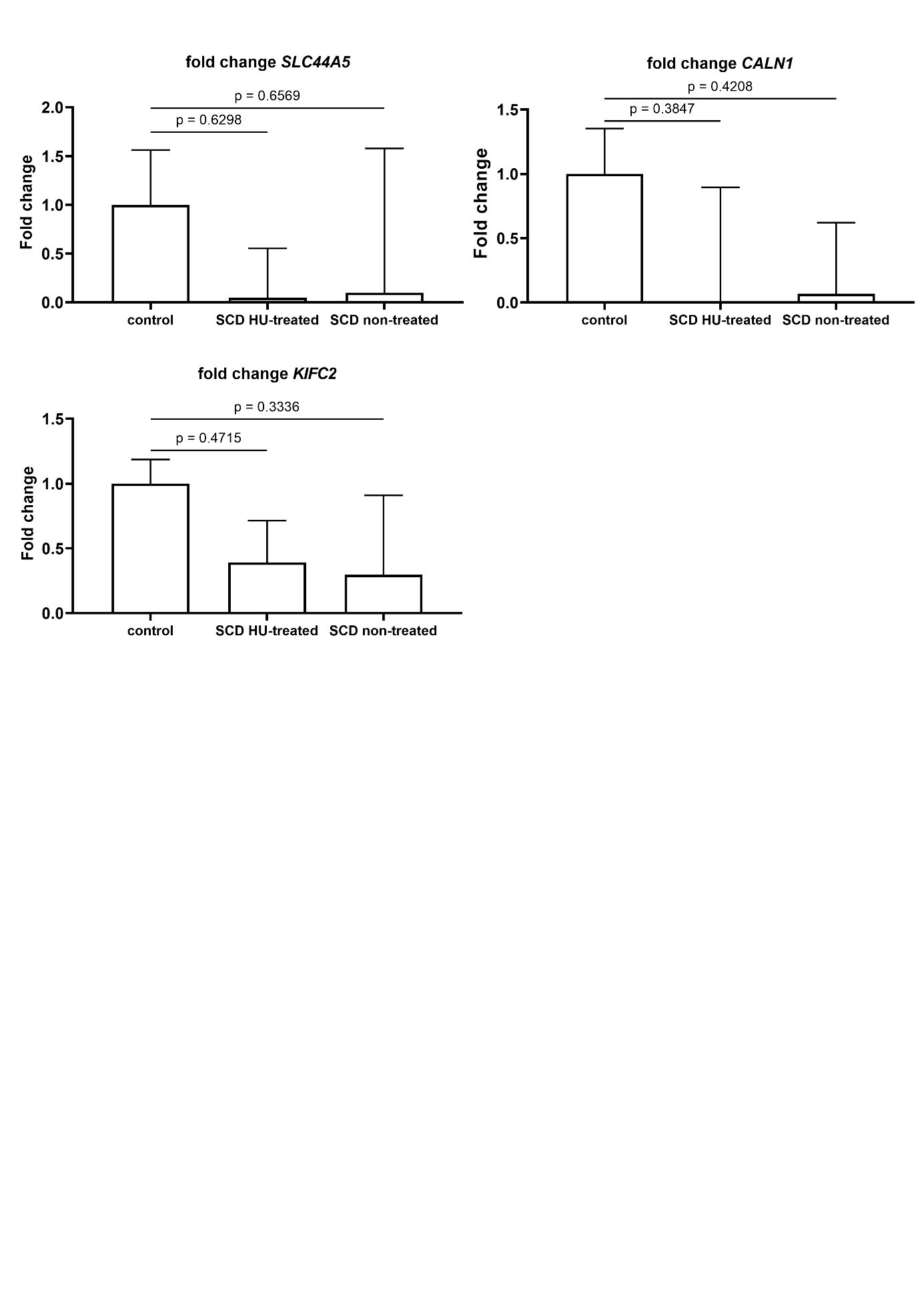
